# A New Detailed Mass Offset Search in MSFragger for Improved Interpretation of Complex PTMs

**DOI:** 10.1101/2025.07.28.667198

**Authors:** Carolina Rojas Ramírez, Fengchao Yu, Daniel A. Polasky, Alexey I. Nesvizhskii

**Affiliations:** Department of Pathology, University of Michigan, Ann Arbor, MI, 48109, USA

## Abstract

Conventional database search methods for proteomics struggle when tasked to identify dozens or hundreds of modifications simultaneously. Open or error-tolerant searches can address this limitation, but at the cost of increased difficulty in downstream interpretation of the results and quantification. We and others have previously described “mass offset” or multi-notch searches that sit in between closed and open searches, allowing simultaneous search for hundreds of modifications with more straightforward downstream interpretation than open search. The original mass offset searches were closer to open search, lacking the ability to restrict modifications to specific amino acids. Here, we describe a new “detailed” mass offset (DMO) search implemented in the MSFragger search engine, which allows each mass offset to have its own site restrictions and fragmentation rules. The benefits of DMO search over existing mass offset searches are shown with three example searches of complex modification sets: nearly one hundred post-translational modifications, fast photochemical oxidation of proteins (FPOP)-derived modifications, and amino acid substitutions. The DMO search further improves the interpretability of results by reducing ambiguity in site localization, particularly when modifications have overlapping masses, and provides benefits that scale with the complexity of the search.

## INTRODUCTION

Post-translational modifications (PTMs) of proteins play essential functional and regulatory roles^1^. One of the major advantages of mass spectrometry-based proteomics is the ability to detect post-translationally (or otherwise) modified peptides based on mass differences, making it perhaps the most important method for analysis of protein post-translational modifications^2^. There are many types of protein PTMs and tremendous diversity and complexity within many types^3^, such glycosylation or epigenetic marks, as well as crosstalk between types^4^. In addition to biological modifications, there is a wide variety of chemical modifications introduced to proteins, either artefactually during sample preparation or deliberately to accomplish specific analyses, such as in chemoproteomics^5^ or protein foorprinting^6^. This vast space of biological and chemical modifications of proteins represents a tremendous challenge for data interpretation tools that match mass spectra to specific peptide sequences.

As a result, several distinct approaches have been developed to search for peptide modifications. The most common method, implemented in nearly all proteomics search engines, is to add copies of a base peptide sequence to the search database for each modified form of the peptide, according to a set of rules defining which modification(s) can be placed on which amino acid(s)^7^. This “variable modification” or “closed” search allows for relatively strict control over the modification search space, as each modification can be restricted to a small number of amino acid sites and most search engines offer control over the maximum number of modifications that can be present on a given peptide. These strict definitions are amenable to downstream processing tools, such as for peptide and protein quantitation, as the modification definitions can readily be passed to the downstream tools. The simplicity and control of this method comes at the cost of a combinatorial expansion of the search space when multiple types of modification are considered. For complex searches the combinatorial expansion becomes prohibitive, as millions of permutations of each base peptide sequence can be generated^8,9^.

For searches of complex modification types, alternative approaches are thus necessary. If the closed search, with specific modification-amino acid pairs pre-defined and encoded in the search database is one extreme, “error-tolerant”^10^ or “open”^11,12^ searches represent the other extreme. In these searches, a large mass difference, or “delta mass” between the mass of the peptide sequence and the observed precursor mass is allowed, so that any modification(s) of the peptide that fall within that range can be identified **(Figure 1A)**. This means that even unexpected or undefined modifications can be found, and any combination of modifications can be identified by their summed mass without needing to enumerate each of the many modification-site permutations. Variations on this theme include tag-based open searching^13,14^ and open spectral library searching^15^. With computational optimization, particularly including fragment ion indexing pioneered by MSFragger, modern open search methods are capable of search speeds comparable to those of non-indexed closed searches despite traversing a much larger search space. Because of their continuous delta mass range, open search results typically require additional processing and interpretation to group observed delta masses together and match each group to a specific modification. While several tools are available to automate this process^16-19^, converting all observed delta masses to clearly defined modifications for downstream quantitation and other processing remains challenging.

**Figure 1.**
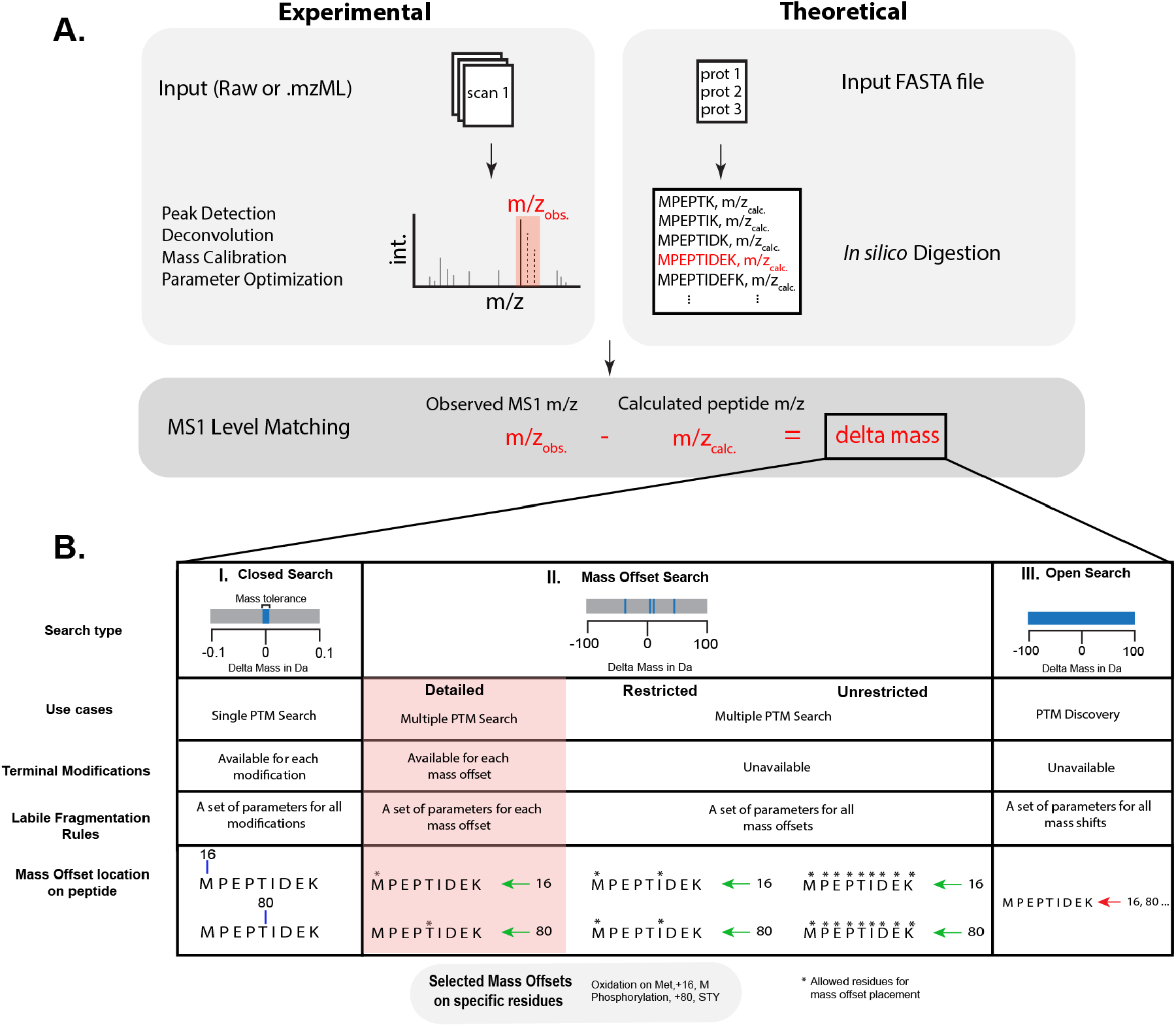
Overview of PTM Searches in MSFragger. **A**. MS1 precursor masses are compared to calculated peptide sequence masses (resulting from *in-silico* protein digestion) to determine the PSM delta mass. **B**. PTM search types available in MSFragger. **I. Closed Search:** Modifications are added directly to the peptide sequence encoded in the peptide index. The delta mass must be within a narrow mass error tolerance (e.g., 20 ppm). Combinations of modifications are allowed, resulting in combinatorial expansion of the search space when multiple modifications are specified. **II. Mass Offset Searches:** Modification masses are not added to peptide sequences and are instead matched as delta mass values within a narrow tolerance. Unrestricted MO search allows any offset to be placed on any site, restricted MO search allows a single set of residues and fragmentation rules for all mass offsets, and detailed MO search allows each offset only at its own defined site(s) and with its own fragmentation rules. **III. Open Search:** Large delta masses are allowed between the peptide sequence and observed mass to match any modification within the mass range with no restriction on allowed sites.

In many cases, neither closed nor open searches fully meet the needs of the analysis, particularly for PTM-enriched data and complex chemical modifications, such as those typical in chemoproteomics and structural proteomics methods. In these searches, the PTM enrichment or known chemical modification(s) mean the most common modifications are known *a priori*, so a fully open search to discover common modification masses is not necessary, and downstream quantitation, challenging for open searches, is often desirable. However, the complexity of expected modifications means closed searches can be difficult to impossible to implement due to the combinatorial search space expansion. “Mass offset”^20^ or “multi-notch”^21^ searches have been developed to provide an alternative in between fully closed and open searches, in which only specific delta masses are allowed, rather than a broad range as in open search. Mass offset searches allow for many rare modifications to be considered without generating an enormous search space, as only a single mass offset is allowed per peptide, preventing any combinatorial expansion. Mass offset searches can also be coupled with variable modifications in a hybrid mode, allowing common modifications to be specified as variable mods (allowing multiple per peptide) and rare modifications as mass offsets to tune the search space.^22^

We have previously implemented a mass offset search mode in the MSFragger^12^ search engine and added several features to it, such as support for labile modifications^23,24^. As originally implemented, this mass offset (MO) search was conceptually quite close to an open search, restricting the mass range to specific allowed offsets, but otherwise providing no restrictions, such as to the amino acid sites where mass offsets were allowed. Here, we describe a new implementation of a “detailed mass offset” (DMO) search, in which each mass offset can be restricted to specific amino acid site(s) and can be provided with specific labile fragmentation rules (**Figure 1B**). This DMO search is thus closer along the spectrum to a closed search, where all modifications are provided with specific sites, but retains the ability to handle extremely complex modification sets by precluding combinatorial expansion of the modifications. We demonstrate the utility of this DMO search in several complex search applications, including extended mass offset searches for hundreds of PTMs, fast photochemical oxidation of proteins (FPOP) structural proteomics, and amino acid substitution searches. In the less complex cases, the primary benefit of the DMO search is in more facile interpretation of the output data and improved localization of modifications compared to open searching. However, for more complex cases, the ability to restrict the search space in precise ways enables substantial improvements in quality and enables searches that were previously prohibitively difficult. The DMO search is available since MSFragger 4.1+ and has been incorporated into several workflows in the FragPipe (https://fragpipe.nesvilab.org/) computational environment.

## METHODS

### Datasets

Three datasets are used to demonstrate the benefits of DMO mode for PTM searches. For the “common PTMs” analysis, data from Chick et al. (PXD001468) was downloaded from PRIDE^25^. Briefly, the data consisted of 24 fractions of tryptic peptides from HEK293 cells^11^. Samples were fractionated using basic pH reverse-phase liquid chromatography, desalted, and then analyzed using a Q-Exactive Orbitrap mass spectrometer (Thermo Scientific, San Jose, CA).

For the amino acid substitutions analysis, two replicate LC-MS/MS proteomics injections of a 1:1 mixture of *Escherichia coli* and *Salmonella typhimurium* peptides were downloaded from MassIVE^26^ (MSV000092029)^27^. After LC separation using a C18 column, data were acquired on a TIMS-TOF Pro instrument (Bruker Daltonics, Bremen, Germany) in DDA-PASEF mode^28^. A list of singly substituted peptides (SSPs) of *S. typhimurium* compared to *E. coli* was produced following the steps as described in the original publication^27^. Briefly, both *E. coli* and *S. typhimurium* proteomes were *in silico* digested by ProteaseGuru^29^ and input into a script called “FindSSP.py” (from the original publication), which produces a target list of peptides that differ by a single amino acid between *E. coli and S. typhimurium*.^27^

For the “FPOP” analysis, data from Espino *et al*. was downloaded from PRIDE (PXD019290)^22^. Briefly, intact *Caenorhabditis elegans* nematodes were oxidized using hydrogen peroxide flash photolyzed with an excimer laser. After protein extraction, tryptic digestion, and clean up, samples were analyzed by LC coupled to a Orbitrap Fusion Lumos Tribrid mass spectrometer (Thermo Scientific, San Jose, CA).

### Data Analysis

Raw data files were converted to mzML^30^ format with vendor peak centroiding and removal of 0-intensity points using MSConvert version 3.0.22340-77d5bfc. Data was analyzed using FragPipe v23.0, including MSFragger (v4.2)^12^ for database search and Philosopher (v5.1.1)^31^ for peptide assignment validation, protein inference, and FDR filtering.

FragPipe configuration files with all parameters used for each search are available in the supporting data (see Data Availability). The Mass-Offset-commonPTMs and FPOP workflows have also been made available as workflow templates in FragPipe. Briefly, all workflows used stricttrypsin as the digestion enzyme, as well as a peptide length between 7 to 50 residues and cysteine carbamidomethylation was set as fixed modification. For the Common PTMs search and Amino Acid Substitution searches, precursor and fragment mass tolerances were set at 10 ppm and 20 ppm, respectively. For the FPOP search, both precursor and fragment mass tolerances were set at 20 ppm. The list of mass offsets used for each search can be found in the supplementing data.

FragPipe was run on a Linux Server (RedHat 8.10) with an Intel Xeon Gold 6354 processor with 36 CPU cores and 2 threads per core, and 768 GB of RAM. Protein FASTA databases were formatted using Philosopher to include common contaminants and an equal number of reversed sequence decoys. All databases were downloaded from UniProt: *C. elegans* (UP000001940), *E. coli* (UP000000625), *S. typhimurium* (UP000001014) and *H. Sapiens* (UP000005640). Results from FragPipe psm.tsv output files were analyzed with Python scripts to assess the efficacy of DMO mode.

### Detailed Mass Offset Search Implementation

For DMO search, mass offsets (masses) with allowed sites and labile fragment ions (if applicable) can be provided via the FragPipe GUI or loaded from a tabular text format. Standard mass offset searches in MSFragger perform a fragment-indexed search for each mass offset window, meaning that fragments from peptide-delta mass combinations that fall within tolerance of each provided mass offset are considered serially. Restricted offset searches apply universal filters (to all mass offsets) at search time to ignore sites and/or ions that do not follow the specified restrictions. To implement amino acid site restrictions and labile fragmentation rules for each mass offset individually in DMO mode, all sites and ions for all mass offsets are initially considered in the search, as in standard MO searches. Mass offset-specific restrictions are applied for search-time filtering to exclude any potential matches that violate the provided site and/or fragmentation rules for mass offset being considered. Final localization is performed analogously to localization-aware open search^20^, except considering only allowed sites (and/or peptide termini) for the matched mass offset.

### Group-based FDR Estimation

When searching complex modification sets with large search spaces, if modified peptides comprise a small minority of all peptide-spectrum matches (PSM)s, the determination of a score threshold for FDR filtering will be dominated by unmodified peptides with a much larger search space. This can lead to overly liberal FDR estimation for modified peptides. To obtain a proper threshold for each PSM type, group-based filtering in Philosopher allows specification of three PSM groups based on their peptide modification status **(Table S1)**: unmodified, defined (common) modifications, and all other (rare) modifications. The “other” category will contain any observed modifications/mass offset that are not included in the “defined” group, and these modifications will override “defined” modification/mass offset if a PSM has both types of modifications/mass offsets.

For FPOP searches, oxidation on residues with medium and high reactivity (MFHILVWY) were defined and all other FPOP-related modifications were sent to the ‘other’ group. For AA substitution searches, oxidation on M was defined, and mass offsets of amino acid substitutions and all other PTMs considered were in the “other” group. Lastly, due to the requirement of inputting mass offset and residue pairs, to apply group-based FDR, when using regular MO search, all possibilities of mass offset/residue pairs needed to be included. While in DMO search, only the allowed mass offset residue pairs were included.

### Data Availability

FragPipe workflow files containing all search parameters used were deposited at https://zenodo.org/records/16537316. FragPipe result .tsv files for each workflow were included in the Zenodo Repository. Lastly, the detailed mass offset lists used for each workflow were also saved in the Zenodo Repository.

## RESULTS

Compared to the original mass offset search, DMO search has the potential to provide two main benefits when searching for modifications with differing amino acid specificities and/or fragmentation behavior. First, restricting the allowed amino acid sites and/or labile fragment ions reduces the search space, as fewer peptides and sites are considered as possible matches to spectra, potentially improving the sensitivity at a given FDR. Second, restricting the amino acid sites prevents unexpected localizations from being reported, simplifying interpretation of the results. To demonstrate these benefits, we compared the results of traditional MO with DMO searches for three analysis types in terms of the number of PSMs identified at 1% FDR and accuracy of localization and identification.

### Dataset 1: Common PTMs

The first analysis is a PTM-discovery search, using mass offsets for nearly 100 PTMs, tested in a fractionated HEK293 cell line dataset previously used to evaluate open search methods^11^. This analysis simulates a common use case for DMO search, as it enhances the main advantage of a mass offset-based PTM discovery workflow relative to open searching, namely, interpretability of the results. To quickly screen data for common PTMs and artefacts, we often use mass offset searches for a large list of common modifications as an alternative to a fully open search. A list of the most common PTMs was gathered from Unimod^32^ and MetaMorpheus^21^. Some of these common PTMs can be localized to cysteine, so the fixed carbamidomethylation modification was subtracted from the mass of the PTM to produce the final mass offset. A total of 95 mass offsets were searched in both MO and DMO searches. While using MO search, all these mass offsets can be placed on any amino acid, resulting in more ambiguous and incorrectly localized PSMs, whereas when using DMO search, the possibilities for mass offset placement are restricted to the expected amino acid type(s) for each modification.

To compare the results of MO and DMO searches in each analysis, PSMs reported by each search were classified into several categories **(Figure 2)**. First, PSMs were categorized by their modification state **(Figure 2A)**. If no modifications (or only fixed modifications) were present, the PSM was considered unmodified. The remaining PSMs were divided based on whether they contained a mass offset, yielding variable modification (only) and mass offset categories. In the second level of categorization, the mass offset-containing PSMs were divided based on the relevance of their match to the PTM represented by the mass offset **(Figure 2B)**. Mass offsets matching the expected localization, e.g., a phosphorylation mass on a S, T, or Y residue, were called “Allowed” **(Figure 2B, inset table)**. If the mass offset is placed on a residue that is not allowed, but the peptide sequence contains allowed residues, the PSM is considered “Mislocalized”. Lastly, if the mass offset is placed in a PSM where the matched peptide sequence does not contain any allowed residues, the PSM was considered “Not Allowed”. The DMO search cannot produce mislocalized or unallowed results, by definition, as each offset is restricted only to the intended sites. Finally, the PSMs that were categorized as “Mislocalized” and “Not Allowed” in regular MO search were tracked (by scan number) to the corresponding PSM in the DMO search **(Figure 2C)**. While MO search initially reported more modified PSMs than DMO search, a closer look at the composition of these PSMs reveals that the DMO search finds more correctly localized PSMs, as many PSMs in the MO search are either “Mislocalized” or, in a few cases, “Not Allowed” **(Figure 2B)**. Comparing the “Mislocalized” PSMs from MO search to the corresponding PSMs in DMO search shows that the majority (61%) become “Allowed”, indicating a simple repositioning of the mass offset to an allowed site in the DMO search, while 25% are “filtered out,” indicating the score of the relocalized form was not good enough to pass FDR filtering. Looking at the “Not Allowed” PSMs from MO search, only 8% become “Allowed” in the DMO search, while 86% are “Filtered out” from the DMO search. The filtered out PSMs are lost from DMO search for several reasons. Some are simply a misinterpretation of an otherwise correct peptide identification, such as a combination of two modifications (or a modification and a precursor monoisotope selection error) being reported instead of the correct modification **(Figure 3)**. Some result from chimeric spectra or an incorrectly recorded precursor charge state, where the correct peptide sequence is identified but with an erroneous delta mass due to a mismatch with the recorded precursor. FragPipe includes Crystal-C^16^, a module created to account for the correction of precursors in chimeric spectra that can reduce these issues, however, it is not currently compatible with saving the mass offsets as variable modifications (as done in the searches presented here to enable detailed comparison between search types). And some are truly incorrect peptide identifications with the wrong peptide sequence due to a combination of previously mentioned factors **(Figure S1)**. The DMO search thus simplifies the interpretation of the results by correctly localizing modifications mislocalized by MO search and removing PSMs with the correct peptide sequence but difficult to interpret modification or precursor recording errors, as well as removing some PSMs with incorrect peptide sequences.

**Figure 2.**
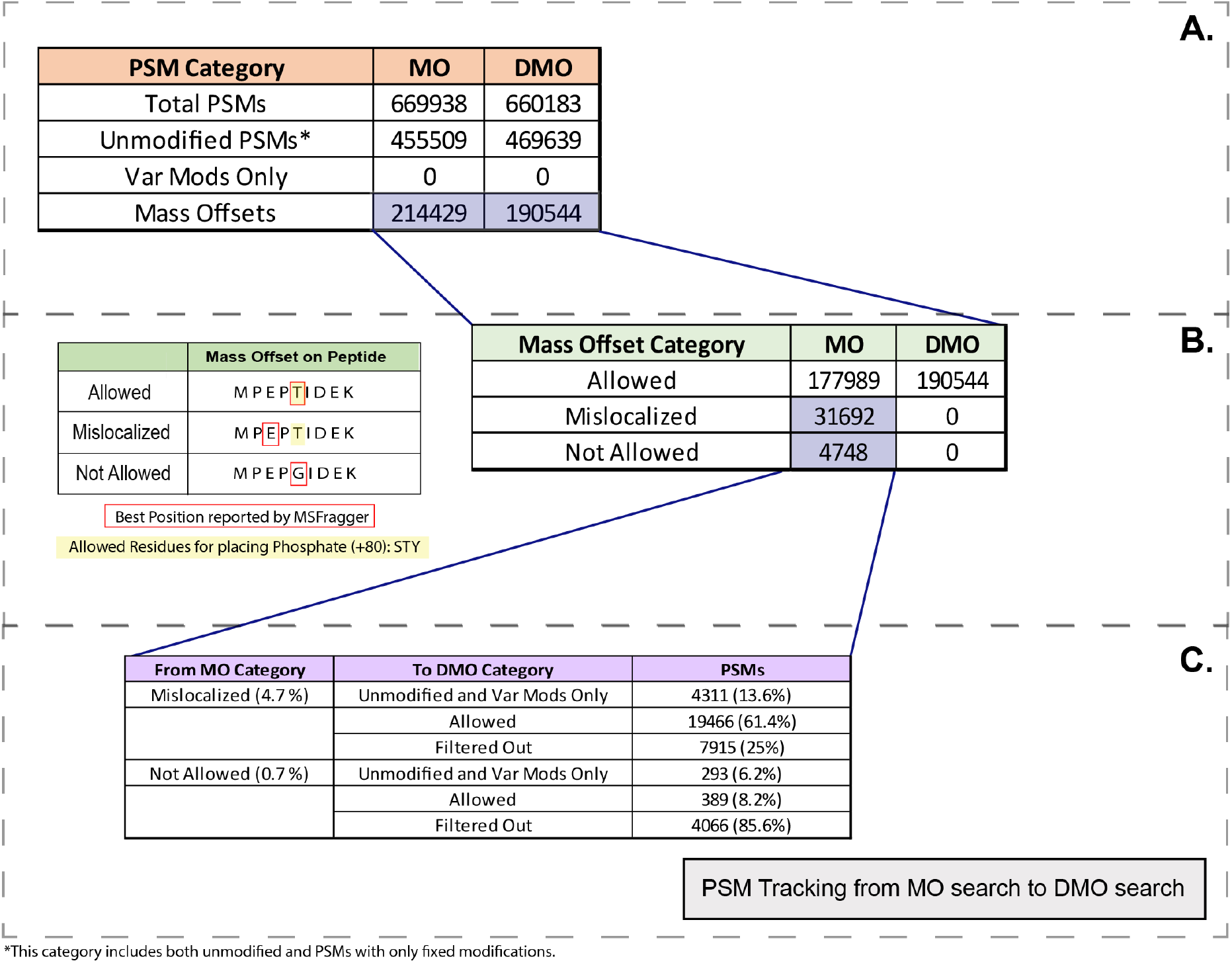
Comparing common PTMs analysis results from MO and DMO searches. **A**. Total PSM counts identified in MO and DMO searches. There were no variable modifications used in this search. **B**. Mass Offset-containing PSMs categorized by the type of mass offset placement on the peptide. The inset table describes the types of placements possible. If the mass offset is placed on an allowed residue (specified by the user), the placement is consider “Allowed”. If the mass offset is placed not on allowed residue but allowed residue(s) are present in the peptide, the PSM is considered a “Mislocalized” placement. If there are no allowed residues in the peptide, the mass offset is considered to have a “Not Allowed placement. DMO only positions mass offsets on allowed residues by definition. **C**. Tracking “Mislocalized” and “Not Allowed” PSMs from MO to DMO search. In the “From MO Category”, the categories names are shown, as well as their total percentage of all PSMs identified in the dataset.

**Figure 3.**
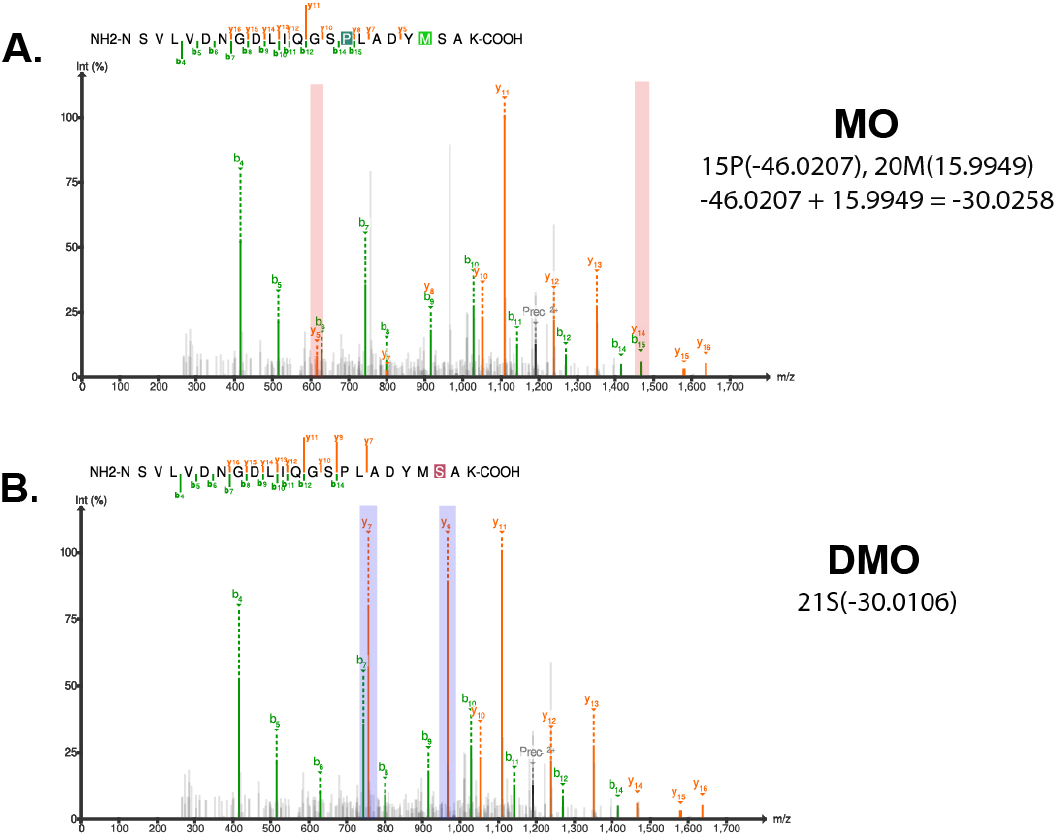
Detailed Mass Offset Improves Peptide Spectrum Matches. **A**. PSM reported by regular Mass Offset (MO) search with both a variable modification and a mass offset with masses that partially cancel. **B**. PSM identified by Detailed Mass Offset search (DMO), matching a single modification with a mass shift of −30 Da, which is the total modification mass found on A (within mass error tolerance). While several low intensity fragments are not matched anymore (highlighted in red rectangles on A.) using DMO search, two high intensity fragments are matched instead (highlighted in blue rectangles on B.).

### Dataset 2: Fast Photochemical Oxidation of Proteins

Next, we analyzed an *in vivo* FPOP dataset previously analyzed by FragPipe^22^, as the photochemical labeling involved produces a very complex search space. In FPOP experiments, single proteins or whole proteomes are exposed to hydroxyl radicals which induces several types of oxidative labels.^34^ While our FPOP search considers only 16 unique modification masses, compared to nearly 100 in the common PTMs analysis, several of these modifications can be found on nearly any amino acid (e.g., oxidation can occur on 19 of the canonical 20 residues) causing a dramatic search space expansion for variable modification searches.^22^ Previously, an FPOP workflow was developed in FragPipe using a hybrid search mode with the most common modifications set as variable modifications and the rest set as mass offsets. The hybrid search was capable of identifying more PSMs with FPOP modifications much faster than a search with only variable modifications.^22^ Here, we assess the impact of the DMO search on the same dataset, comparing to the hybrid MO search previously performed.^22^

As in the common PTMs analysis, the regular MO search identified more mass offset-containing PSMs than the DMO search. However, only half of the mass offset-containing PSMs in the MO search are placed on expected sites (“Allowed”) (**Figure 4B**). The majority of the non-allowed PSMs are due to mislocalization, though a significant number of PSMs fall in the not allowed category as well. The combination of many variable modifications and the fact that many of the provided mass offsets are expected only on a single or few amino acids poses a significant challenge for the regular MO search. When tracking the “Mislocalized” and “Not Allowed” PSMs, from MO search to the DMO search, 45% and 94% of PSMs were “Filtered out”, respectively (**Figure 4C**). As seen previously with the common PTMs analysis, many of the “Mislocalized” PSMs can be rescued to the correct localization by DMO search, while most of the unallowed PSMs are removed. The DMO search thus simplifies the interpretation of the results by correctly localizing modifications not localized properly by MO search. PSMs with the correct peptide sequence but difficult to interpret modifications or with incorrect peptide sequences are filtered out from the results when using DMO search.

**Figure 4.**
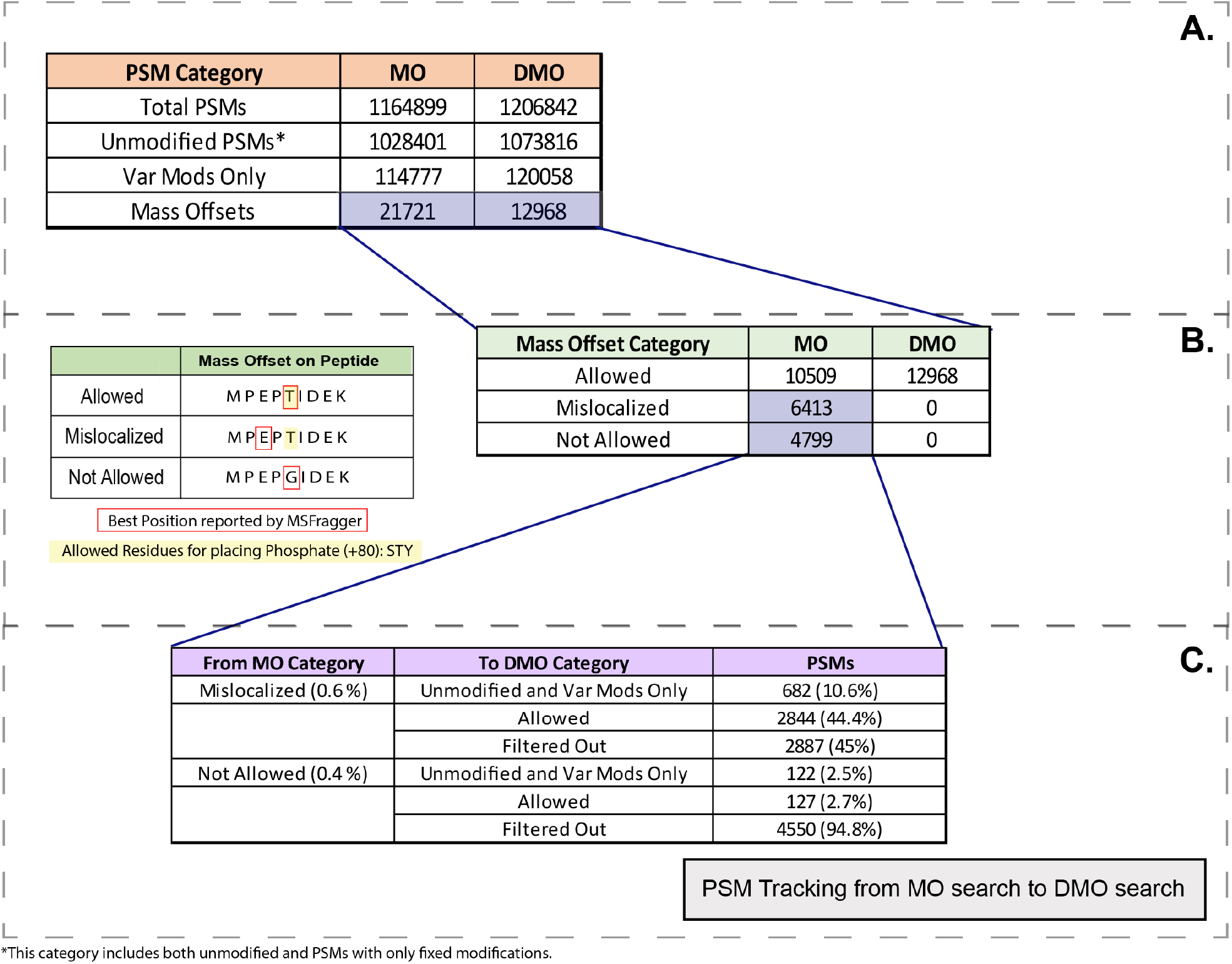
Comparing FPOP analysis results from MO and DMO searches. **A**. PSM totals by modification type. **B**. Mass Offset-containing PSMs are categorized by the type of Mass Offset placement on the peptide, as in figure 2. **C**. Tracking PSMs with “Mislocalized” or “Not Allowed” mass offset placements from MO to DMO search.

### Dataset 3: Amino Acid Substitutions

Lastly, we selected a search for single amino acid substitutions, as this is an emerging area of interest for proteomics searches. Incorporation of unexpected residues, based on the genome-define protein sequence, can lead to detrimental effects on protein function^35^. Characterizing amino acid substitutions directly in MS proteomics data has the potential for untargeted identification of substitutions throughout the proteome, but requires advanced search strategies. In particular, we chose a dataset which provides a ground-truth positive control for the evaluation of AA substitution identification ^27^ by mixing two closely related species, in this case *E. coli* and *S. typhimurium*, and analyzing their peptides together. The combination results in a dataset containing homologous peptides, including many that differ by a single residue, producing a set of known singly substituted peptides (SSPs).

Substitution searches are typically intractable with conventional proteomics search methods due to the large number of possible modifications when considering swapping any of 20 amino acids to any other. Previously, open searches have been used to detect substitutions^36^, however, analyzing open search results is not straightforward, because many substitution mass differences overlap with each other and those of common PTMs and artefactual modifications^37^. Searching for all possible amino acid substitutions with MO search can improve search speed and reliability vs open search, but localization and interpretation remain challenging. Compared to previous methods using open or MO search results, here we show that the DMO search provides improved accuracy from reducing the localization space for substitutions with similar masses.

The detailed mass offset list was constructed by calculating the mass differences for each possible amino acid substitution using a Python script. Residues I and L were considered an interchangeable residue due to their equal mass. Also, X to C or C to X (X is any residue except cysteine) substitutions were corrected for carbamidomethylation, which was set as a fixed modification on cysteine. The resulting mass offset list contained 308 unique masses. Several common PTMs were included in the search due to having the same mass difference as one or more AA substitution(s) to avoid incorrectly calling modified peptides as substituted. As most of these modifications were rare, only methionine oxidation was included in the defined modification set for group FDR, with all other modifications and substitutions in the rare set. As observed in each Common PTMs and FPOP analysis, in the regular MO search there is a slightly higher number of identified mass offset-containing PSMs **(Figure 5A)**. However, unlike the previous analyses, 91% of PSMs were “Allowed” in MO search **(Figure 5B)**. This is partly due to the high degree of overlap in substitution masses, as most peptides will have several residues that are allowed for a given mass, reducing the number of PSMs categorized as “mislocalized” or “not allowed” based on the input rules. When tracking the “Mislocalized” and “Not Allowed” PSMs, from MO search to the DMO search, 51% and 89.5% of PSMs were “Filtered out”, respectively, similar to the other analyses (**Figure 5C**). This emphasizes that DMO search improves the interpretability of the results by filtering out most PSMs that are not allowed and rescuing mislocalized PSMs to the correct site.

**Figure 5.**
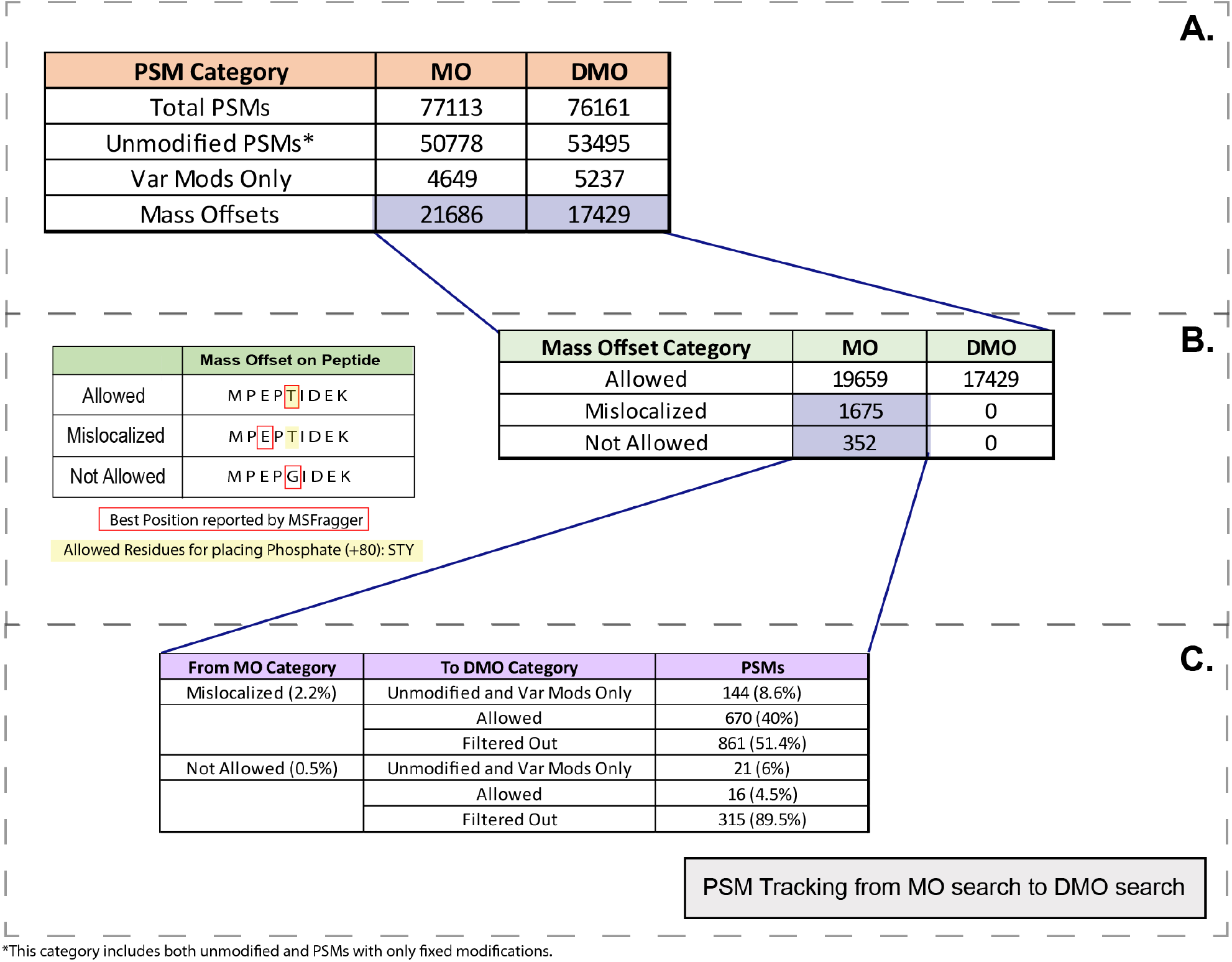
Comparing Amino Acid Substitutions analysis results from MO and DMO searches. **A**. PSM totals by modification type. **B**. Mass Offset-containing PSMs are categorized by the type of Mass Offset placement on the peptide as in figure 2. **C**. Tracking PSMs with “Mislocalized” or “Not Allowed” mass offset placements from MO to DMO search.

Finally, we assessed the MO and DMO searches using the known singly substituted peptides (“SSPs”) resulting from *S. typhimurium* peptides that differ from *E. coli* peptides by a single amino acid. When searching using a database containing only one species, in this case *E. coli*, and the list of amino acid substitution modifications, *S. typhimurium* SSPs can be identified as *E. coli* peptides with an amino acid substitution and compared to the ground truth *S. typhimurium* peptide to determine the accuracy of the amino acid substitution identification.

In **Figure 6A**, a flowchart shows the procedure to determine if an *E. coli* mass offset-containing PSM is in fact a *S. typhimurium* SSP. First, not all *E. coli* peptides have a single AA substitution counterpart in *S. typhimurium*, so only those peptide sequences of *E*.*coli* with a counterpart were considered as possible SSP peptides. Then, any mass offset and residue pair that matched any common (non-substitution) PTM was removed to focus only on substitutions. If the substitution mass offset and residue pair led to the expected *S. typhimurium* counterpart sequence, it was considered a SSP PSM. In **Figure 6B**, the breakdown of PSMs following the flowchart on **Figure 6A** is shown. While there are more mass offset-containing PSMs identified in MO search, there were 849 more SSPs identified in the DMO search. This is largely due to mislocalization of putative substitution masses in the MO search, as any change in localization changes which AA substitution is being called. Additionally, near-equal mass substitutions (within the 20 ppm precursor mass tolerance employed) and off-by-1 precursor monoisotope picking errors more easily confound the MO search given its much larger localization space for most substitution masses. The restrictions of the DMO search alleviate these issues and allow for more correct SSPs to be reported. However, even in the DMO search, there are several substitution masses that are shared between numerous amino acid pairs, so while the DMO search provides improved accuracy compared to the MO search (which in turn is more reliable than fully open search), additional validation of potential substitutions remains essential.

**Figure 6.**
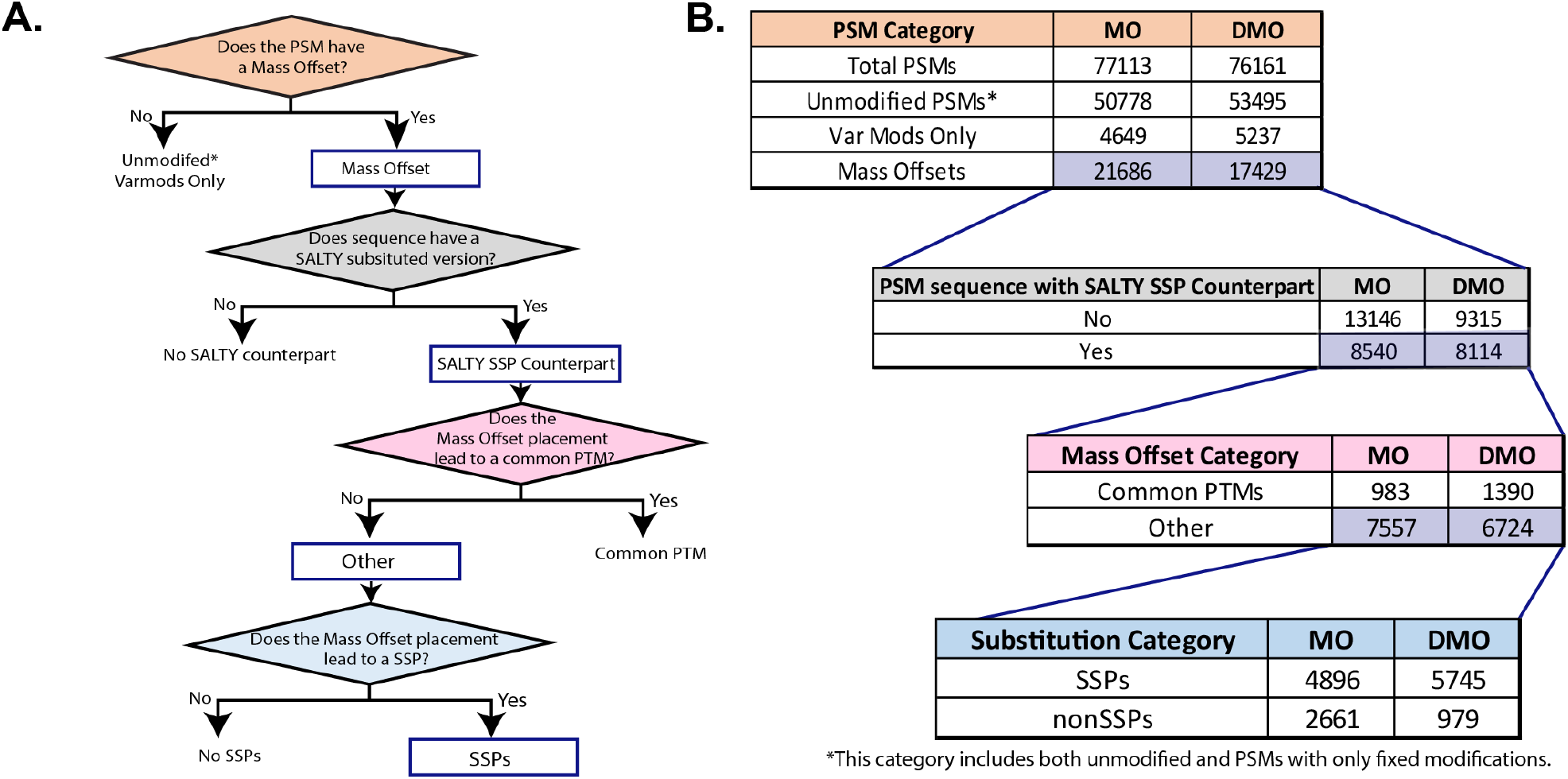
MO and DMO results compared to ground truth substitution peptides. **A)** Flowchart of the process to determine the successful identification of a ground truth SSP. The color of each decision rhombus matches the color of the table header in B. Only mass offset containing PSMs are considered in this analysis. If the PSM sequence has a known SSP counterpart in the *S. typhimurium* proteome, the mass offset placement will be further analyzed to see if it produced a substituted peptide or not. **B)** Breakdown from PSMs to SSPs and comparison between both MO and DMO searches. Not all mass offset-containing PSMs have a *S. typhimurium* SSP to compare in the ground truth set. The mass offset-containing PSMs that do are further analyzed to determine if the mass offset and its placement produced a *S. typhimurium* SSP.

## CONCLUSIONS

Traditional closed searches for modified peptides suffer from a combinatorial search space expansion that precludes simultaneous searches for many modifications. Meanwhile, open searches prevent combinatorial search space expansion, but sacrifice straightforward interpretation of the data, as an offset in a peptide might indicate one or more than one modification is present. In addition, due to the complexity of searching for tens or hundreds of modifications at once, chimeric spectra, low quality spectra, high abundant unknown modifications, and suboptimal mass accuracy during data acquisition can make it difficult for the search engine to determine an appropriate precursor that leads to an accurate delta mass. Combining these factors with a mass offset search with unrestrictive placements can lead to modified PSMs with unexpected mass offsets, sequences or both. In this study, we introduced an in-between option with the detailed mass offset search mode in MSFragger, allowing one mass offset per identified peptide and only allowed at specific locations. DMO mode prevents a large expansion of the search space, and the results are easier to interpret than open or unrestricted MO searches. We demonstrated the simplified interpretation of DMO results in three analyses of complex modification sets. When using DMO in very complex searches, total PSM counts were similar to MO search but with a slight decrease in the number of mass offset-containing PSMs. The vast majority of these lost PSMs are in the “Not Allowed” mass offset localization category. For example, when using regular MO mode for amino acid substitution search, 65% of all possible SSPs were identified, compared to 85% identified when using DMO. As improvements in mass accuracy and resolution continue to expand the palette of PTMs and other modifications accessible to proteomics, DMO search is well positioned to support increasingly complex modification searches and provide straightforward and reliable interpretation of the results. The DMO search mode is now available MSFragger 4.1+ and FragPipe v22+.

## Author Contributions

D.A.P. and A.I.N.: conceptualization; C.R.R, D.A.P., and F.Y.: method development; C.R.R and D.A.P.: writing original draft; C.R.R., D.A.P., F.Y. and A.I.N.: writing, review and editing; A.I.N.: supervision; A.I.N.: funding acquisition.

## Acknowledgements

This work was funded in part by the National Institutes of Health grants R01-GM-094231 and U24-CA271037.

