## Supplementary Information for "A New Detailed Mass Offset Search in MSFragger for Improved Interpretation of Complex PTMs"

| **Table of Contents** | **S1** |
|---|---|
| **Table S1**. PSM groups formed by Philosopher for Group-based FDR. | **S2** |
| **Figure S1.** Precursor Charge State Mismatched and Chimeric Spectrum Example. | **S3** |

**Table S1. PSM groups formed by Philosopher for Group-based FDR.** Fixed modifications are treated as unmodified.

| Group | PSM Modification Types |
|---|---|
| Unmodified | Unmodified |
| Defined | Defined Modification |
| Other | Other Modifications |
| | Other Modifications + Defined Modification |

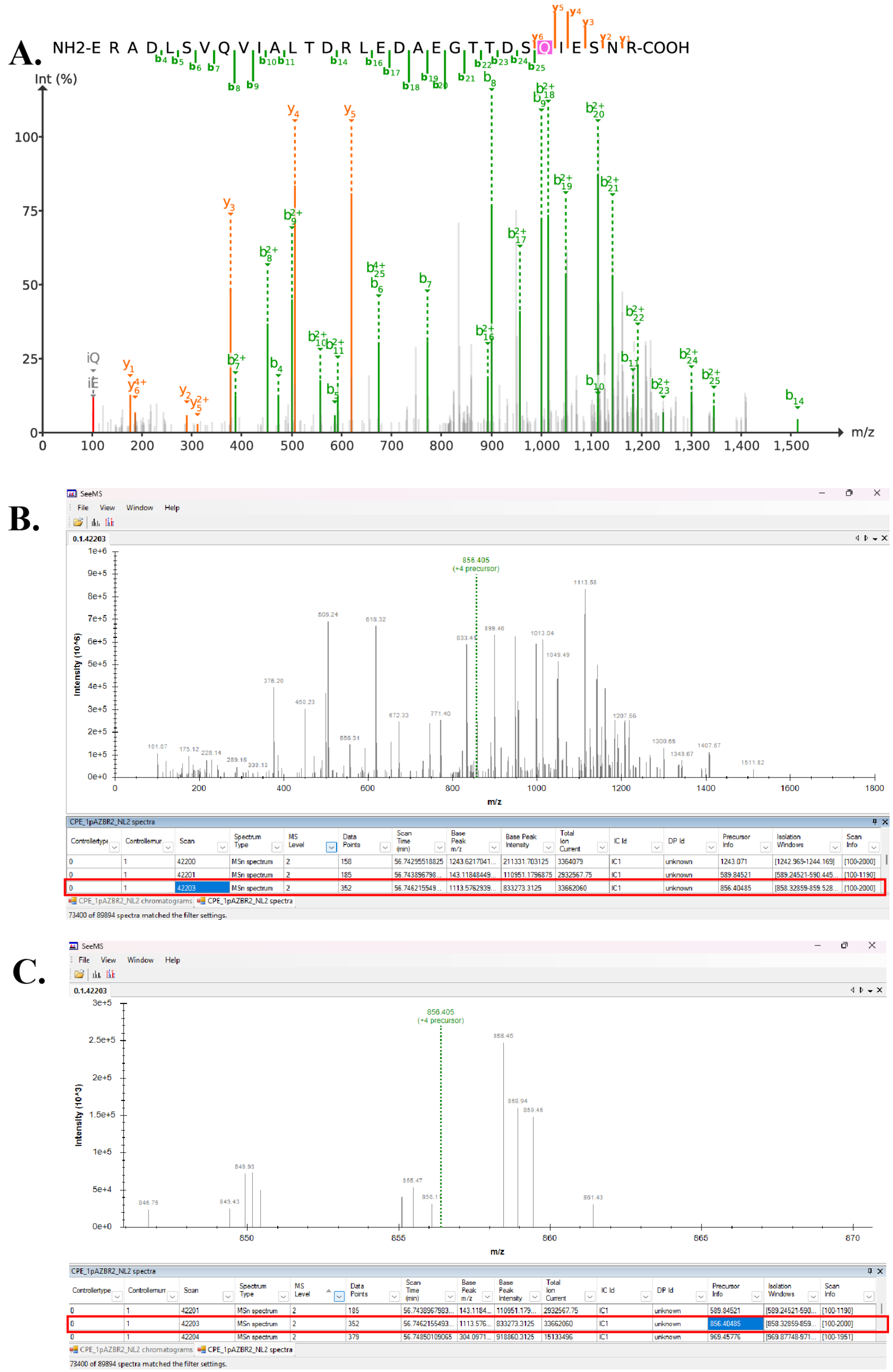


**Figure S2. Precursor Charge State Mismatched and Chimeric Spectrum Example. A.** PSMs (1pAZ_sample_BR2_NL2-42203.4) from FPOP dataset where the mass offset (-9.0368) it is not placed on an allowed residue. Due to a wrong precursor selection, the mass offset is not correct. **B.** Precursor selection error. Reported precursor (m/z 856.4085, +4) peak was not properly selected as it is not within the isolation window (m/z 858.3 – 859.5). Scan highlighted red rectangle. **C.** Precursor Charge State Error. The reported precursor at m/z 856.4 not only has the wrong m/z but also charge. Zooming in, the precursor possibilities are m/z 858.45 with z = 1 and z = 2. Two overlapping precursors might explain the appearance of what seems to be two peptide fragmentation distributions on A. Scan highlighted red rectangle. PSM was filtered out DMO search results.
